# Novel inhibition mechanism of SARS-CoV-2 main protease by ebselen and its derivatives

**DOI:** 10.1101/2021.03.11.434764

**Authors:** Kangsa Amporndanai, Xiaoli Meng, Weijuan Shang, Zhenmig Jin, Yao Zhao, Zihe Rao, Zhi-Jie Liu, Haitao Yang, Leike Zhang, Paul M. O’Neill, S. Samar Hasnain

**Affiliations:** Molecular Biophysics Group, Department of Biochemistry and System Biology, Institute of System, Molecular and Integrative Biology, Faculty of Health and Life Sciences, University of Liverpool, Liverpool, L69 7ZB, United Kingdom; Department of Molecular and Clinical Pharmacology, Institute of Translational Medicine, Faculty of Health and Life Sciences, University of Liverpool, Liverpool, L69 3BX, United Kingdom; State Key Laboratory of Virology, Wuhan Institute of Virology, Chinese Academy of Sciences, Wuhan, Hubei, 430071, China; Shanghai Institute for Advanced Immunochemical Studies and School of Life Science and Technology, ShanghaiTech University, Shanghai 201210, China; Department of Chemistry, Faculty of Science and Engineering, University of Liverpool, Liverpool, L69 7ZD, United Kingdom

**Keywords:** SARS-CoV-2, COVID-19, Main protease, Ebselen, Drug development

## Abstract

The global emergence of SARS-CoV-2 has triggered numerous efforts to develop therapeutic options for COVID-19 pandemic. The main protease of SARS-CoV-2 (M^pro^), which is a critical enzyme for transcription and replication of SARS-CoV-2, is a key target for therapeutic development against COVID-19. An organoselenium drug called ebselen has recently been demonstrated to have strong inhibition against M^pro^ and antiviral activity but its molecular mode of action is unknown preventing further development. We have examined the binding modes of ebselen and its derivative in M^pro^ via high resolution co-crystallography and investigated their chemical reactivity via mass spectrometry. Stronger M^pro^ inhibition than ebselen and potent ability to rescue infected cells were observed for a number of ebselen derivatives. A free selenium atom bound with cysteine 145 of M^pro^ catalytic dyad has been revealed by crystallographic studies of M^pro^ with ebselen and MR6-31-2 suggesting hydrolysis of the enzyme bound organoselenium covalent adduct, formation of a phenolic by-product is confirmed by mass spectrometry. The target engagement of these compounds with an unprecedented mechanism of SARS-CoV-2 M^pro^ inhibition suggests wider therapeutic applications of organo-selenium compounds in SARS-CoV-2 and other zoonotic *beta*-corona viruses.

## Introduction

The recent emergence of severe acute respiratory syndrome coronavirus 2 (SARS-CoV-2) has resulted in a global pandemic of corona virus disease 2019 (COVID-19) with confirmed infection cases of over 70 million and 1.5 million fatalities within a year. SARS-CoV-2 is the most devastating zoonotic coronavirus to infect humans following SARS-CoV-1 and MERS-CoV (Middle East respiratory syndrome) which emerged in 2002 and 2012, respectively^1^. Similar to the other coronaviruses, SARS-CoV-2 primarily infects the respiratory system and develops critical pneumonia that highly necessitates ventilatory support and intensive care, particularly in elderly and immunocompromised individuals^2^. Whilst there have been tremendous strides forward in the development of vaccines, the current roll-out is supply and time limited. Several vaccine candidates have been developed in advance stage and recently approved for mass immunity first in the UK^3^. However, some effective vaccines need to be stored at deep freeze temperatures that may not be deployable in developing areas of the world. Moreover, some mutations in SARS-CoV-2 genome may impact the effectiveness of developing vaccines to control the virus^4,5^. These underline the requirement for the parallel development of therapeutic options for SARS-CoV-2 treatment.

SARS-CoV-2 is an enveloped, positive-sense, single-stranded RNA virus with a large genome of about 30,000 nucleotides. The whole genome of SARS-CoV-2 is 96% identical to a bat coronavirus and closely related to SARS-CoV-1 with 80% sequence identity^6^. Two overlapping polyproteins, pp1a and pp1ab, are encoded by the replicase gene (ORF 1a/1b) that constitutes two-thirds of the genome. The remainder of the genome encodes for accessory and structural proteins such as the spike glycoprotein, envelope protein, matrix protein and the nucleocapsid phosphoprotein^7^. pp1a and pp1ab are proteolytically digested into 15 non-structural proteins (NSPs) by the two viral proteases. The 33.8kDa main protease (M^pro^) or NSP5 is responsible for cleaving polyproteins at 11 cleavage sites giving NSP4-9 and NSP12-15. The released NSPs form the viral RNA polymerase complex are involved in replication and transcription of fresh virus in the host. Due to vital function of in SARS-CoV-2 life cycle and absence of homologous proteins in human, M^pro^ has been extensively explored by high-throughput screening of re-purposed druggable compounds^8^ and fragments^9^ to devise effective inhibitors aimed at arresting the growth of SARS-CoV-2 in host’s cell.

SARS-CoV-2 M^pro^ is a homodimeric enzyme consisting of three domains^8^. The substrate-binding site with a catalytic dyad of His41 and Cys145 is located between chymotrypsin-like domains I and picornavirus 3C protease-like domain II. Domain III plays an important role in M^pro^ dimerization through a salt-bridge interaction between protomers. Several inhibitors and fragments have been co-crystallised and identified to block catalytic cavity^8–10^.

Ebselen is an organo-selenium molecule that can function as a glutathione peroxidase and peroxiredoxin mimic^11^. It has been shown to form a selenosulphide bond with thiol groups of cysteine (Cys) on a number of proteins which results in anti-inflammatory, anti-microbial and neuroprotective effects^12–14^. Moreover, ebselen is being investigated in clinical trials as a potential therapy for stroke, hearing loss and bipolar disorder with good safety profiles with no adverse effects^15–17^. Recently, ebselen was identified in high-throughput screen as a potential hit of SARS-CoV-2 M^pro^ inhibitor. Molecular dynamics simulations suggested that ebselen is able to bind at two probable sites^18^. One is at Cys145 within the catalytic cavity through a selenosulphide bond, and another is at the dimerization region. However, no experimental data for the site of its binding in SARS-CoV-2 M^pro^ has become available.

In our previous work, we have designed CNS penetrant ebselen-based derivatives and demonstrated their good neuroprotective effects and low cytotoxicity in cell-based and mouse models of motor neuron disease^19^. Here, ebselen and five derivatives were assessed for their inhibition of SARS-CoV-2 M^pro^ and anti-coronaviral activity. Two of these ebselen-based selenium compounds exhibit greater inhibitory effectiveness than ebselen against M^pro^ enzyme and SARS-CoV-2 replication. We show from co-crystallographic studies of M^pro^ enzyme with ebselen and another potent compound (MR6-31-2) that these compounds solely bind at the M^pro^ catalytic site by donating a selenium atom, forming a covalent bond and blocking the histidine-cysteine catalytic dyad. We propose that the ebselen-enzyme drug protein adduct is hydrolysed by the conserved water in the catalytic pocket. The release of phenol by-product has been confirmed by mass spectrometry studies of M^pro^ incubated with compounds. This novel unpreceded mechanism of inhibition and direct observation of covalent binding of the selenium atom together with sub-micromolar antiviral activity provides a rational for utilising ebselen as potential therapy and improving selenium-based compounds using the ebselen scaffold for greater anti-coronaviral activity.

## Results

### M^pro^ enzymatic and antiviral activities of ebselen and derivatives

In our previous study, ebselen and some selenium-based derivatives have were developed as neuroprotective agents in relation to motor neuron disease^19^. The co-crystalised structures of those compounds with superoxide dismutase 1 (SOD1) were proven to form a selenylsulphide bond with Cys111 at dimer interface. Thus, ebselen and derivatives were considered for their reactivity with Cys145 and their potential for impairing proteolytic activity of M^pro^ in an attempt to arrest the growth of SARS-CoV-2. A fluorescence resonance energy transfer assay was conducted to evaluate the inhibition level against M^pro^ enzyme of ebselen and five other selenium-based derivatives. Fig. 1 shows chemical structures, inhibitory curves against M^pro^ and the half-maximal inhibitory concentrations (IC_50_s) for each of the compounds. The data clearly demonstrates that these compounds including ebselen are potent M^pro^ inhibitor with sub-micromolar levels of IC_50_. Some compounds are twice as effective for M^pro^ inhibition than the parent ebselen, especially MR6-7-2 and MR6-18-2. All of these compounds were also assessed for *in vitro* antiviral activity against SARS-CoV-2 infected human Vero cells. All of the compounds tested were somewhat superior to ebselen with MR6-31-2 being nearly three times more effective with an EC_50_ of 1.8 μM (Fig. 1g). These results indicate clear on-target interaction of these compounds with M^pro^ with significant inhibitory power for SARS-CoV-2 and as such potential for development as treatments for COVID-19 patients.

**Fig. 1:**
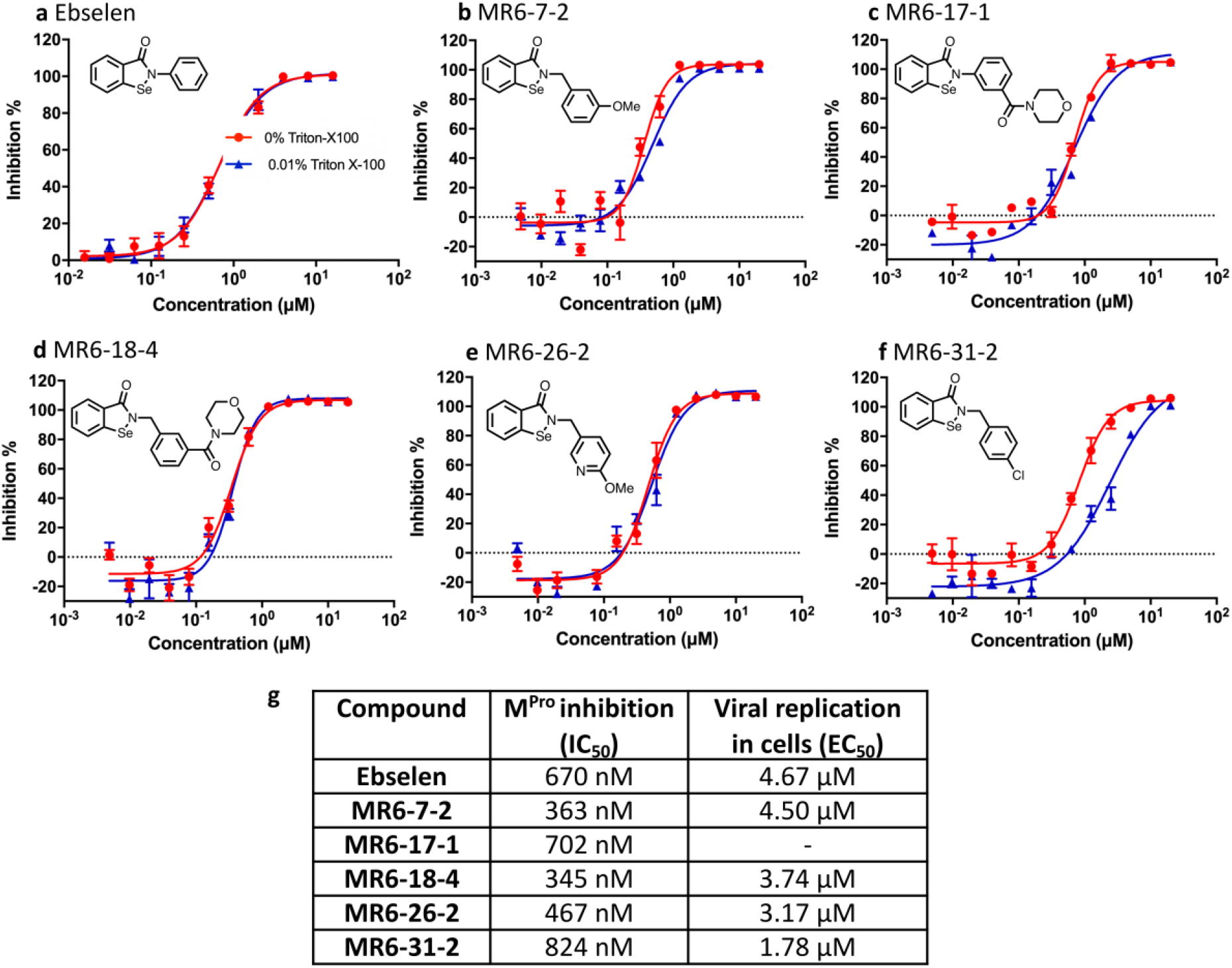
Chemical structures, *in vitro* M^pro^ inhibition and cell-based antiviral assays of ebselen and five derivatives. *In vitro* M^pro^ inhibitory curves of **a**, ebselen, **b**, MR6-7-2, **c**, MR6-17-1, **d**, MR6-18-4, **e**, MR6-26-2, and **f**, MR6-31-2. **g**, IC_50_s of M^pro^ inhibition and EC_50_s of viral replication in human cells.

### Structures M^pro^ with ebselen and MR6-31-2 reveal selenium atom bound in catalytic site

The interaction of ebselen and its derivative with M^pro^ was directly visualised by co-crystallisation of organoselenium compounds with M^pro^. The structures of ligand-free and M^pro^ complexes with ebselen and MR6-31-2 have been solved at the resolution of 1.6-2.0Å. The statistics of data collection and structure refinement is summarised in Table 1 for ligand-free M^pro^, M^pro^-ebselen and M^pro^-MR6-31-2. There is only one M^pro^ protomer found in asymmetric unit where the global structures are almost identical (Fig. 2a). The M^pro^ catalytic site including cysteine-histidine dyad of individual structures are given in Fig. 2b-c. Electron density is clearly visible and allowing amino acid residues and water molecules to be defined accurately. Interestingly, a clear patch of electron density is observed between Cys145 and His41 in co-crystallised crystals of M^pro^-ebselen and M^pro^-MR6-31-2. This density is too strong to be a water, but the size is too small for the corresponding complete inhibitors. To identify the origin of this clear density, anomalous electron density map of selenium was calculated from diffraction data using X-ray at the wavelength near selenium absorption edge (0.97Å). No selenium anomalous density is observed in ligand-free enzyme (Fig. 2b), but a strong density is present only at the cysteine-histidine catalytic dyad in ligand-treated M^pro^ structures (Fig. 2c and 2d). Thus, selenium atom was modelled into the density with the distances of 2.2 Å away from Cys145 and His41. Calculated B-factor suggest that the occupancy of selenium in M^pro^-ebselen and M^pro^-MR6-31-2 crystals are 60% and 80%, respectively. These data are consistent with ebselen and MR6-31-2 primarily binding at M^pro^ catalytic pocket and form a selenyl-sulphide bond with Cys145. There was no density associated with the organic backbone of either ebselen or MR6-31-2 in the co-crystallographic structures. As only selenium atom is observed in the enzyme’s active site, it is the inactivation of cysteine by selenium that results in the inhibition of M^pro^ activity and viral replication. Interestingly, selenium binding does not affect the conformation of surrounding amino acid residues within the active site. According to crystallographic evidence, we noted that ebselen and selenium-based derivatives have unusual mode of action by selenation of M^pro^ catalytic dyad.

**Table 1.**
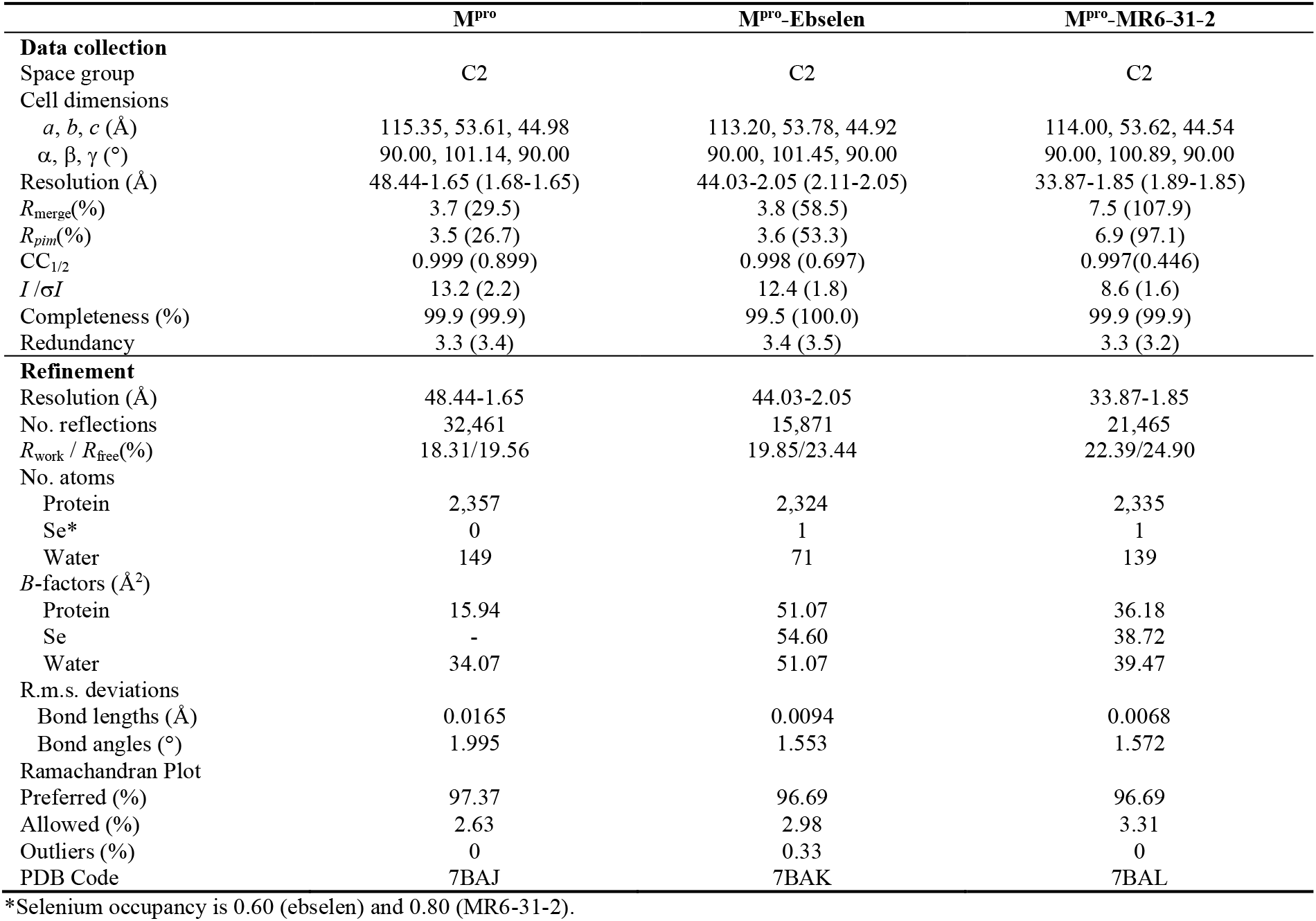
Crystallographic data collection and refinement statistics of M^pro^ and the complexes with ebselen and MR6-31-2.

**Fig. 2:**
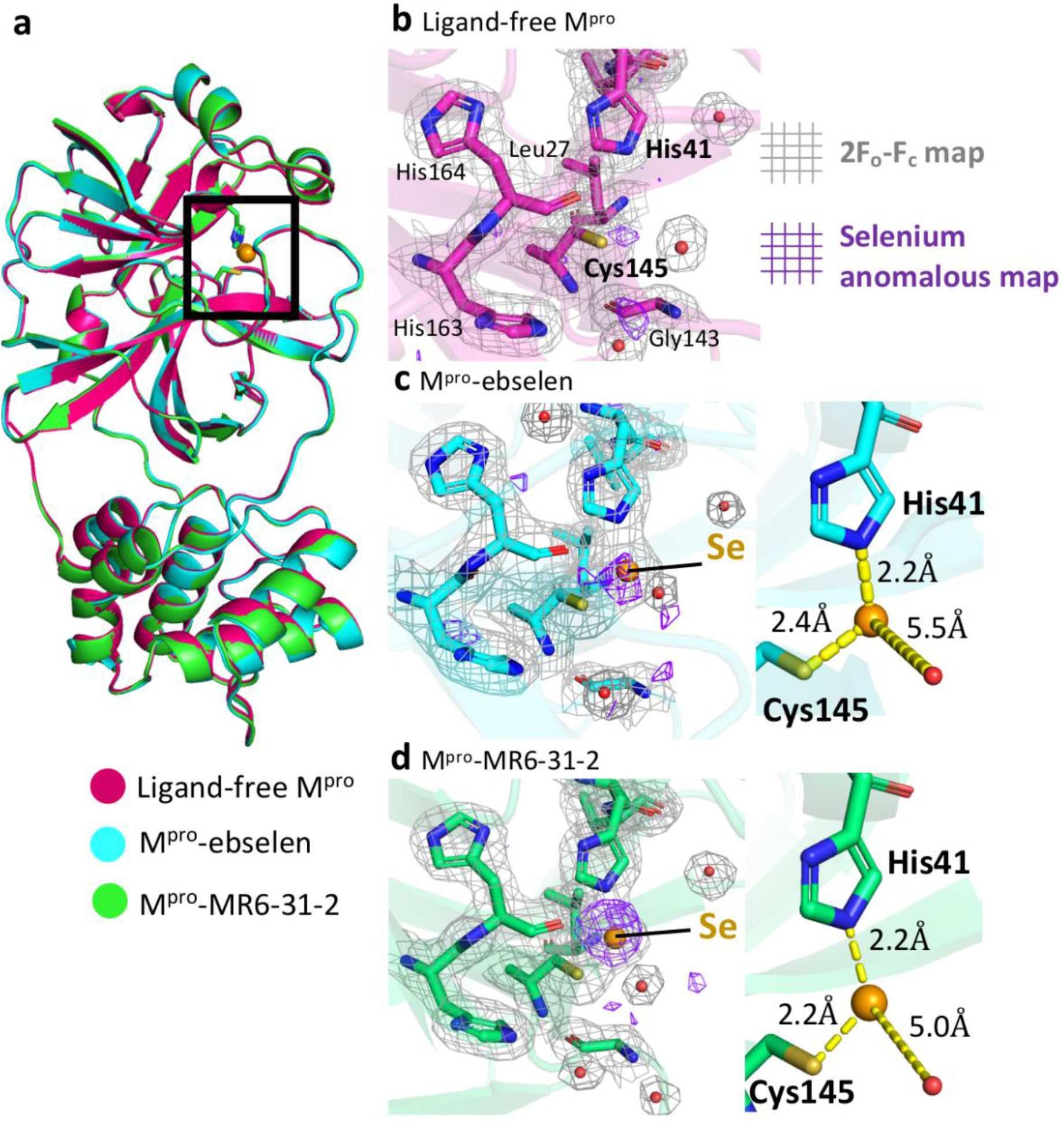
Crystallographic structures of ligand-free M^pro^ and the complexes with ebselen and MR6-31-2. **a,** Cartoon representation of superimposed structures of ligand-free M^pro^ (magenta), M^pro^-ebselen (cyan) and M^pro^-MR6-31-2 (green). The M^pro^ catalytic site is highlighted in a black box. Close-up views of catalytic site of **b,** ligand-free M^pro^, **c,** M^pro^-ebselen, and **d,** M^pro^-MR6-31-2. Electron density (2F_o_-F_c_) map is shown as grey mesh at 1σ. Anomalous signal of selenium is shown as purple mesh at 3σ. Selenium atom and waters are shown as orange and red spheres, respectively

### LC-MS characterization of salicylanilide generated by hydrolysis of ebselen

Co-crystalised structures of M^pro^ with ebselen and MR6-31-2 demonstrated that the compounds would inhibit by selenation at Cys145 of catalytic dyad. This evidence suggests that the compounds may be hydrolysed within M^pro^ active site that releases its phenolic by-product (salicyanilide for ebselen, Fig. 3a). In order to identify the hydrolysis product derived from M^pro^-ebselen adduct, LC/MRM-MS method was optimized using standard salicyanilide. A representative chromatogram from standard salicyanilide (85.2 ng/mL) is shown in Fig. 3a. Samples obtained from the incubation of ebselen showed strong peaks at 6.37 minutes corresponding to salicylanilide (Fig. 3b). The MS/MS spectrum of molecular ion at m/z 214 showed product ions at m/z 121 and m/z 94, which are attributed to the ions derived from the cleavage of the amide bond (Fig. 3c). To measure the levels of salicyanilide formed in the incubation, an 8-point calibration line was generated for salicylanilide in BSA (r^2^ = 0.99, data not shown). The measured concentration of salicylanilide in these samples after 240 minutes was 1.28 ng/ml. The formation of salicyanilide in the incubation of M^pro^ with ebselen is time dependent (Fig. 3d).

**Fig. 3:**
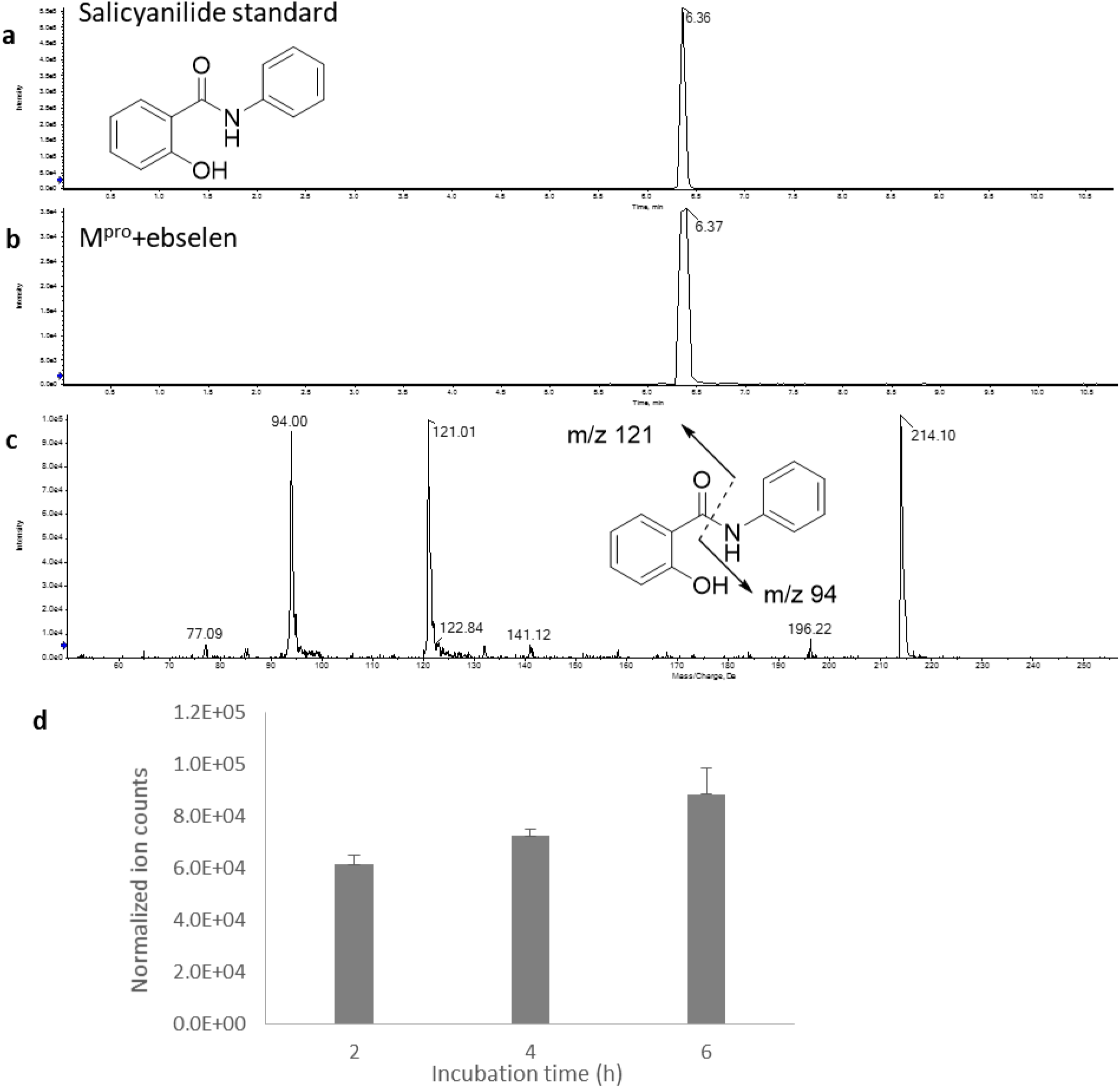
Representative chromatograph for salicyanilide standard and its formation in the incubation of M^pro^ with ebselen. **a,** The peak at 6.36 minutes retention time corresponds to salicyanilide standard (85.2 ng/mL). **b,** Salicyanilide was detected in the incubation of M^pro^ with ebselen. **c,** MS/MS spectrum shows the characteristic fragments derived from salicyanilide. **d,** Time-course of salicyanilide formation.

## Discussion

From a chemical mechanism of action perspective, we fully expected to see the SARS-CoV-2 M^pro^ drug-adduct **2** from ebselen **1**, through nucleophilic attack of the Cys145 thiolate on the electrophilic selenium centre as shown in Fig. 4. Unlike other M^pro^ covalent inhibitors^8,20,21^, the organic framework of ebselen was not present in the co-crystallographic structures with evidence of extrusion of selenium from the ebselen core in an unprecedented reaction at a cysteine protease active site. We propose that His41 can assist a water mediated attack on intermediate adduct **2** in an S_N_Ar type hydrolysis reaction with intermediate **3** possibly stabilised in a manner akin to peptide hydrolysis tetrahedral intermediates within the oxyanion hole of the active site. With increased activity in the drug-design field in the covalent modification of catalytic and non-catalytic thiols, there have been several reports of aromatic warheads tuned with leaving groups (halides for example) to enable nucleophilic aromatic substitution, so the S_N_Ar aspect of the proposed mechanism is with precedent^22–24^. Based on this mechanism, we would expect to see the generation of the hydrolysis product **4**. Using LCMS analysis of the SARS-CoV-2 M^pro^ and comparison with a commercial of **4**, we were able to show that **4** is generated from ebselen by the enzyme in a time-dependent manner. This provides strong support for our proposed mechanism for selenation of the SARS-CoV-2 M^pro^ active site.

**Fig. 4:**
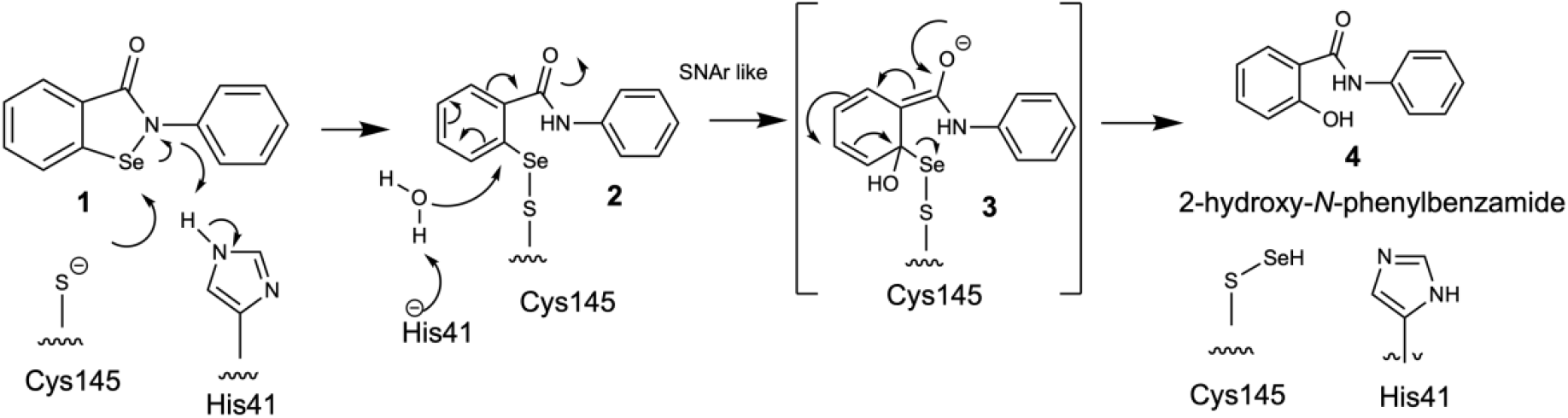
Chemical mechanism for an unprecedented selenation of a cysteine protease enzyme by ebselen.

We have succeeded in obtaining co-crystallographic structure of M^pro^ grown with ebselen and its derivative MR6-31-2, showing selenium coordinates directly to Cys145 upon hydrolysis of the organoselenium compounds. The clear target engagement paves the way for further development for more effective delivery to the catalytic cysteine and greater inhibition whilst having an acceptable safety profile. Though our study is of clear immediate interest for SARS-CoV-2, it has wider therapeutic applications of organo-selenium compounds by novel chemical mechanism of the selenation of cysteines of proteases in other current zoonotic beta coronaviruses and those that may emerge in the future.

## Methods

### Synthesis of compounds

Ebselen was obtained from a commercial supplier (Sigma Aldrich). Other lead compounds were produced and purified at Department of Chemistry, University of Liverpool. Details of the synthesis of ebselen-based derivatives have been described in our previous report^19^.

### Recombinant SARS-CoV-2 M^pro^ production

SARS-CoV-2 M^pro^ gene (GenBank: MN908947.3, residues 3258-3569) containing modified human rhinovirus 3C protease (HRV-3C) cleavage site and 6xHis tag (SGVTFQGPHHHHHH) at C-terminal was cloned into pGEX-6P-1 vector at BamHI and XhoI sites using gene synthesis and cloning services (GenScript, USA). The plasmid was transformed into *E. Coli.* strain BL21(DE3) and cultured at 37°C in 2xYT broth until optical density at 600nm reaches 0.8. M^pro^ expression was induced by the addition of 0.5mM isopropyl ß-d-1-thiogalactopyranoside (IPTG) followed by incubation at 37°C for 5 hours. The bacteria pellet was harvested by centrifugation at 5,000xg, 4°C for 20 minutes and then re-suspended in lysis buffer (20mM Tris pH 7.8, 150mM NaCl) before sonicated on ice. Cell lysate was collected by centrifugation at 30,000xg, 4°C for 30 minutes and then loaded onto a 5mL NiNTA affinity column (HisTrap HP, GE Healthcare) pre-equilibrated with the lysis buffer. M^pro^ bound to NiNTA resin was washed with 100mL of 5mM imidazole in lysis buffer and then eluted with a linear gradient of imidazole from 5mM to 500mM in lysis buffer, 100mL. The fractions containing M^pro^ were pooled together, mixed with recombinant His-tag HRV-3C, and dialysed against 20mM Tris pH 7.8, 150mM NaCl, 1mM DTT at 4°C overnight. The mixture containing M^pro^ was re-loaded through fresh NiNTA resin to remove uncleaved protein and HRV-3C. The His-tag cleaved M^pro^ in the flow-through was buffer-exchanged to 20mM Tris pH 8 using Amicon Ultra centrifugal filter (MWCO. 10kDa, Merck) and then loaded onto 5mL Q Sepharose column (HiTrap Q HP, GE Healthcare). The column was eluted with 100mL of a linear gradient from 0mM to 500mM NaCl in 20mM Tris pH 8. The fractions containing pure M^pro^ were buffer-exchanged to 20mM Tris pH 7.8, 150mM NaCl for activity assay or crystallisation or 25mM ammonium bicarbonate pH 7.5 for LC-MS. The concentration of M^pro^ was determined by ultraviolent absorption at 280nm using a molar extinction coefficient of 32,890 M^−1^cm^−1^.

### M^pro^ inhibition activity assay

The inhibition activity assays were performed using 0.2 μM M^pro^, 20 μM substrate and serial-diluted tested inhibitors in 60 μL reaction buffer consisting of 50 mM Tris-HCl (pH 7.3), 1 mM EDTA. Firstly, M^pro^ was incubated with testing inhibitors at 30 °C for 15 min in reaction buffer. The reaction was then initiated by the addition of a FRET-based peptide substrate Mca–AVLQ↓SGFR-K(Dnp)K (GL Biochem), using wavelengths of 320 nm and 405 nm for excitation and emission, respectively. Fluorescence intensity was monitored with an EnVision multimode plate reader (Perkin Elmer). Initial rate was obtained using the data from the first 10 min by linear regression. To exclude inhibitors possibly acting as aggregators, a detergent-based control was performed by adding 0.01% freshly made-up Triton X-100 to the reaction at the same time. The IC_50_ was calculated by plotting the inhibition rate against various concentrations of testing inhibitor by using a four parameters dose-response curve in GraphPad Prism software. All experiments were performed in triplicate.

### Antiviral activity assay

A clinical isolate of SARS-CoV-2 (nCoV-2019BetaCoV/Wuhan/WIV04/2019) was propagated in Vero E6 cells, and viral titer was determined as described previously^25^. For the antiviral assay, pre-seeded Vero E6 cells (5×10^4^ cells/well) were pre-treated with the different concentration of compound for 1 h and the virus was subsequently added (MOI of 0.01) to allow infection for 1 h. At 24 h post infection, the cell supernatant was collected and vRNA in supernatant was subjected to qRT-PCR analysis as described previously^25^. For cytotoxicity assays, pre-seeded Vero E6 cells were treated with appropriate concentrations of compound. After 24 h, the relative numbers of surviving cells were measured by the CCK8 (Beyotime, China) assay in accordance with the manufacturer’s instructions. All experiments were performed in triplicate, and all the infection experiments were performed at biosafety level-3 (BSL-3).

### Crystallisation and structure determination

Ebselen and other compounds were prepared as 250mM stocks in DMSO. 0.1mM purified M^pro^ was incubated with 1mM compound at 4°C overnight before concentrated to 5-15mg/mL protein. Hanging crystallisation drops were set by mixing of 3 μL of M^pro^, 2.4 μL of reservoir solution (200mM ammonium chloride, 5%glycerol and 16-20% polyethylene glycol (PEG) molecular weight 3350) and 0.6 μL of 1/2560 diluted micro-seed stock. Micro-seed stock was prepared by crushing M^pro^ crystals obtained from an initial hit (well A9 of JCSG+ screen: 200mM NH_4_Cl, 20% PEG3350, Molecular Dimensions) with glass seed beads (Hampton Research). The crystallisation drops were placed against 300 μL corresponding reservoir at 19°C allowing vapour diffusion. Plate crystals of M^pro^ appeared among precipitation within a week. The crystals were cryo-protected in 25% glycerol in reservoir solution before snap-frozen in liquid nitrogen. X-ray diffraction experiments were carried out at 100K using 0.9795Å beam on I04 beamline of Diamond Light Source, UK. Identification of selenium in active site was made by anomalous x-ray diffraction measurement in the same crystal using 0.9795Å wavelength at 3.0Å resolution. The data was integrated by using DIALS^26^ and scaled by using Aimless^27^. Phase problem was solved by molecular replacement with MOLREP^28^ using a SARS-CoV-2 M^pro^ structure (PDB: 6Y2E) as an initial model. Structure models were edited manually in COOT^29^ and refined by using Refmac5^30^. Geometry and quality of final models were validated by using MolProbity^31^.

### Liquid chromatography-mass spectrometry (LC-MS)

10mg/mL M^pro^ in 25mM ammonium bicarbonate pH 7.5 (20 μL) was incubated with 1mM ebselen at 37°C for 0, 2, and 4 h. At the end of incubation, 2.5mM acetaminophen (10 μL) was added as an internal standard to normalize extraction. Then, loading and compounds of interest were extracted by adding ice-cold acetone (250 μL). Standard curve was constructed by spiking salicylanilide (concentration range: 0.1-0.8 μM) into 10mg/mL bovine serum albumin (BSA) solution. After centrifugation at 14000 rpm for 20 min, the extracts were transferred to clean tubes and evaporated in a Speed Vac and reconstituted in 50 μL 30% ACN/0.1% formic acid. 10 μL samples and standards were analysed immediately by a QTRAP 5500 mass spectrometer (AB Sciex,) coupled with an Ultimate 3000 HPLC system (Dionex, ThermoScientific) and a Kinetex C18 column (2.6uM, C18, 50mm × 2.1mm, Phenomenex, Maclesfield, Cheshire, UK). The MS experiments were conducted using electrospray ionization with positive ion detection. A gradient program of acetonitrile (5% for 1 min; 5% to 95% over 5 min; 95% for 2 min; 95% to 5% over 0.1 min; 5% for 4 min) in 0.1% formic acid (v/v) was applied at a flow rate of 300 μL/min. The multiple reaction monitoring transitions for each analyte were as following: salicylanilide 214.2/121.1 and 214.2/95; acetaminophen, 152.1/108.1; other MS parameters, such as voltage potential and collision energy were optimized to achieve great sensitivity. Data acquisition and quantification were performed using Analyst 1.5 software and Multi-Quant 3.0 (AB Sciex).

## Acknowledgement

This work was supported by the BBSRC’s.IAA award 168064. Some of the developments arose from ALSA grant (WA1128). We acknowledge the University’s silver command team and the institute’s technical team for making arrangements so that experimental work could be undertake despite COVID-19 restrictions. We would like to acknowledge the support of the staffs and management of the Diamond Light Source (Didcot, United Kingdom) for the beamtimes and operations at the facility (proposal: mx27113). We thank to Michael Rogers for synthesis of ebselen and derivatives.

## Author contributions

SSH and PMO conceived the work. KA purified protein and performed crystallographic experiment. XM performed mass spectrometry experiment. HY and LZ laboratories performed the enzymatic inhibition and antiviral assay in biosafety level 3. All authors read and approved the final version of the manuscript.

